# Protease target prediction via matrix factorization

**DOI:** 10.1101/275024

**Authors:** Simone Marini, Francesca Vitali, Sara Rampazzi, Andrea Demartini, Tatsuya Akutsu

## Abstract

**Motivation:** Protein cleavage is an important cellular event, involved in a myriad of processes, from apoptosis to immune response. Bioinformatics provides in silico tools, such as machine learning-based models, to guide target discovery. State-of-the-art models have a scope limited to specific protease families (such as Caspases), and do not explicitly include biological or medical knowledge (such as the hierarchical protein domain similarity, or gene-gene interactions). To fill this gap, we present a novel approach for protease target prediction based on data integration.

**Results:** By representing protease-protein target information in the form of relational matrices, we design a model that: (a) is general, i.e., not limited to a single protease family; and (b) leverages on the available knowledge, managing extremely sparse data from heterogeneous data sources, including primary sequence, pathways, domains, and interactions from nine databases. When compared to other algorithms on test data, our approach provides a better performance even for models specifically focusing on a single protease family.

**Availability:** https://gitlab.com/smarini/MaDDA/ (Matlab code and utilized data.)

**Contact:**, or

## 1 Introduction

Protein cleavage is a pivotal process in cell metabolism of both cellular and extracellular matrix. Among other processes, protein cleavage is involved in cell differentiation and cycle control, stress and immune response, removal of abnormally folded proteins and cell death (Rawlings *et al*., 2016). The proteins responsible for cleavage, i.e. the proteases, account for about 2% of all gene products (James, 1999). As a consequence, wrongly regulated proteolytic activity may result in diseases (Rawlings *et al*., 2016; Wang *et al*., 2014). Caspase activity, for example, play a role in Alzhemier’s Disease (Chu *et al*., 2015; Zhao *et al*., 2016), Autoimmune lymphoproliferative syndrome (Lenardo *et al*., 1999), and cancer (Oh *et al*., 2010; Hosgood *et al*., 2008; Lee *et al*., 2009). Currently, several computational methods have been proposed to tackle the key-lock machinery of the protease-protein target recognition (Song *et al*., 2010; Wang *et al*., 2014; Boyd *et al*., 2005; Song *et al*., 2012; Wilkins *et al*., 1999; Barkan *et al*., 2010). Cleavage target models, for example, aim at extracting sequence patterns or frequency matrices from known protein target structures/primary sequences in order to predict the likely cleavage points for proteases to act. Design of such algorithms goes back to more than a decade, confirming protease target prediction as a main research focus in Bioinformatics.

A basic but effective approach to predict the targets of a certain protease consists of a simple BLAST sequence search (Wang *et al*., 2014). The BLAST approach relies on the following assumption: if it is known that a protease *P*_*p*_cleaves a protein target *P*_*X*_, then the more a candidate target *P*_*y*_ shares primary sequence similarity to *P*_*X*_, the more likely *P*_*y*_ will be a target as well. Therefore, in order to infer new targets, researchers can score candidate protein *P*_y_ against its similarity (i.e., the BLAST eValue) with known protease *P*_*p*_ target proteins, [*P*_*X*_, …,*P*_*X*_].However, even if there is a strong BLAST similarity between known target of *P*_*p*_, and the query protein *P*_y_, this does not guarantee that *P*_y_ will be a target of *P*_*p*_ as well. In fact, the cleavage mechanism does not depend on general protein similarity, but to the similarity of specific primary structure patches, influenced by sequential and spatial patterns of specific amino acids in key positions (Song *et al*., 2012; James, 1999). In other words, even if two proteins share a very high sequence similarity overall, the cleavage dichotomy between target and non-target might be due by a difference between only few amino acids pivotal positions of the sequences. For example, the peptidase thrombin cleaves the target if an arginine is present next to the scissile bond, in N-terminal direction, but only if an aspartate or a glutamate is *not* juxtaposed to it in the carboxyl-terminal direction. The pioneering efforts in cleavage target prediction were based on the analysis of primary sequence only, for example with PeptideCutter (Wilkins *et al*., 1999) and PoPS (Boyd *et al*., 2005). The peculiarity of these methods, differentiating them from generic protein interaction prediction algorithms (Marini *et al*., 2011; Planas-Iglesias *et al*., 2013), is the embedding of specific protease knowledge. For example, PoPS searches for candidate cleavage spots through the primary sequence with a short sliding window, considering both cleavage-specific position-specific scoring matrix (PSSM) and cleavage-specific weight vectors. On the other hand, PeptideCutter predicts cleavage sites by applying enzyme-specific rules exploiting amino acid-specific cleavage probability tables. More recently, novel machine learning techniques have been applied to solve the cleavage target prediction problem. These approaches are mostly based on support vector machines (SVMs) and trained on complex protein features, such as structural and physicochemical features, including solvent accessibility or disordered regions (Song *et al*., 2012; Barkan *et al*., 2010; Song *et al*., 2010; Wang *et al*., 2014). Despite improving the quality of the predictions, these recent algorithms still suffer from two major limitations. First of all, they are specific for single proteases, i.e. each SVM model is trained to predict the cleavage protein target of a specific protease. While the known human proteases are hundreds (Rawlings *et al*., 2016), prediction algorithms end up focusing on the most known ones, such as the ones belonging to the Caspase family, or the HIV-1 protease (Singh and Su, 2016). Therefore, these models are protease-specific, and lack of generalization, i.e. a different model needs to be trained for each protease. This led to a well-known bias toward “superstar” proteases, leaving most of the other proteases orphan of adequate prediction models. Note that problem encompasses not just proteases-protein target prediction, but Proteomics prediction in general (Orlowski *et al*., 2007; Lam *et al*., 2016). Furthermore, state-of-the-art algorithms do not leverage on the implicit knowledge crosslinks available from biological and medical ontologies and databases, even if trained on fine-grained structural information. For example, Cascleave2 (Wang *et al*., 2014) exploits Gene Ontology (Gene Ontology Consortium, 2015), InterPro (Finn *et al*., 2017), and KEGG (Kanehisa *et al*., 2017), respectively for GO term, domain, and pathway information. However, these data are elaborated and converted into numerical attributes describing single samples, and then fed to SVMs. With this traditional feature encoding approach, other *indirect*, although relevant, knowledge is not embedded in the prediction model. Examples of ontological relationships not encoded as features are: the domain interactions; the hierarchical relationships between domains and genes; or the overlapping of the same gene across different pathways. Intuitively, it is possible to numerically relate the instances of different data sources with *interacts-with* (e.g. protein *P*_*x*_ interacts with protein *P*_*y*_) or *is-part-of* (e.g. domain *D* is found in protein *P*_*x*_) type of relations, as it happens, for example, in the hierarchically structured Gene Ontology. Biological knowledge-bases are indeed saturated with relational knowledge, and yet this knowledge is not exploited in a typical feature encoding. For example, it is possible to map domains belonging (or not) to proteins from InterPro, and to represent the relations with binary numbers. Similarly, is possible to track protein interactions from STRING (Szklarczyk *et al*., 2017), and represent the relations with continuous scalars in the [0, 1] interval, reflecting the STRING confidence score.

Here we present a novel approach to predict protease-to-protein target interactions overcoming these limits and provide a general model for protease target prediction. Note that our model does not predict the cleavage sites, but scores the probability of a protein to be a feasible target for a given protease. Our approach is based on the representation of different data sources through matrices allowing to directly integrate ontologies and other knowledge sources in the learning model. In other words, our approach accounts for multiple, heterogeneous data sources, without altering the knowledge data structure. For cleavage prediction we exploited the repositories MEROPS, STRING, BLAST similarity, Interpro, Domine (Raghavachari *et al*., 2008), 3did (Mosca *et al*., 2014), UNIProt, BioGRID (Chatraryamontri *et al*., 2017) and KEGG. In the field of recommendation systems, matrix decomposition-based methods demonstrated their power in the prediction of disease subtypes alignment (Gligorijević *et al*., 2016) or drug repositioning (Vitali *et al*., 2016). In this work we apply an data integration method based on non-negative matrix tri-factorization (Žitnik and Zupan, 2015) to predict protein targets for human proteases. Our approach is a *general-purpose model*, and the results confirmed its ability in providing predictions not limited to specific, well-studied proteases. We further showed the proposed method outperforms five existing approaches in terms of both range of application and results (ROC area).

## 2 Methods

The algorithm we utilized is a variant of the non-negative matrix tri-factorization data integration approach (Žitnik and Zupan, 2015; Gligorijević *et al*., 2016; Vitali *et al*., 2016), the main difference consisting in cross-validating the training set in order to find the optimal parameter combination. Relations between proteases, protein targets, genes, pathways and domains, harvested from knowledge databases, are first represented by different matrices. All the resulting matrices are then combined into a single block matrix. An iterative algorithm, guided by relational constraints, decomposes the original matrix into three smaller ones. Novel relations (i.e. novel protease-protein targets) are finally inferred by comparing the original and the reconstructed matrices.

### 2.1 Feature encoding by relational matrices

The basic assumption of tri-factorization is that every data instance (i.e. a relation) is represented by the value of a relational matrix cell. Here, rows and columns are sets of data types, such as proteases, target proteins, or genes. For example, given a set of *N* proteases encoded by *M* genes, the fact that a protease *P*_*n*_ is encoded by gene *G*_*m*_ is indicated by the value “1” of the cell corresponding to *P*_*n*_ row and *G*_*m*_ column of a *N×M* matrix. In this work, we considered five data types: *proteases, protein targets, genes, domains* and *pathways*. All the relationships between the elements of these data types are represented through two types of relational matrices: the *input matrices,* and the *constraint* matrices *θ*_*ij*_ The *input matrices R*_*ij*_ ∊ *R*^*n*_*i*_ × *n*_*i*_^, represent different data types relations (e.g. protease-gene); while the *constraint matrices θ*_*i*_ ∊ *R*^*n*_*i*_ × *n*_*i*_^ represent same data types relations(e.gi. protease-protease). Figure 1 depicts the specific *R* and *θ* matrices utilized in this work.

**Fig. 1.**
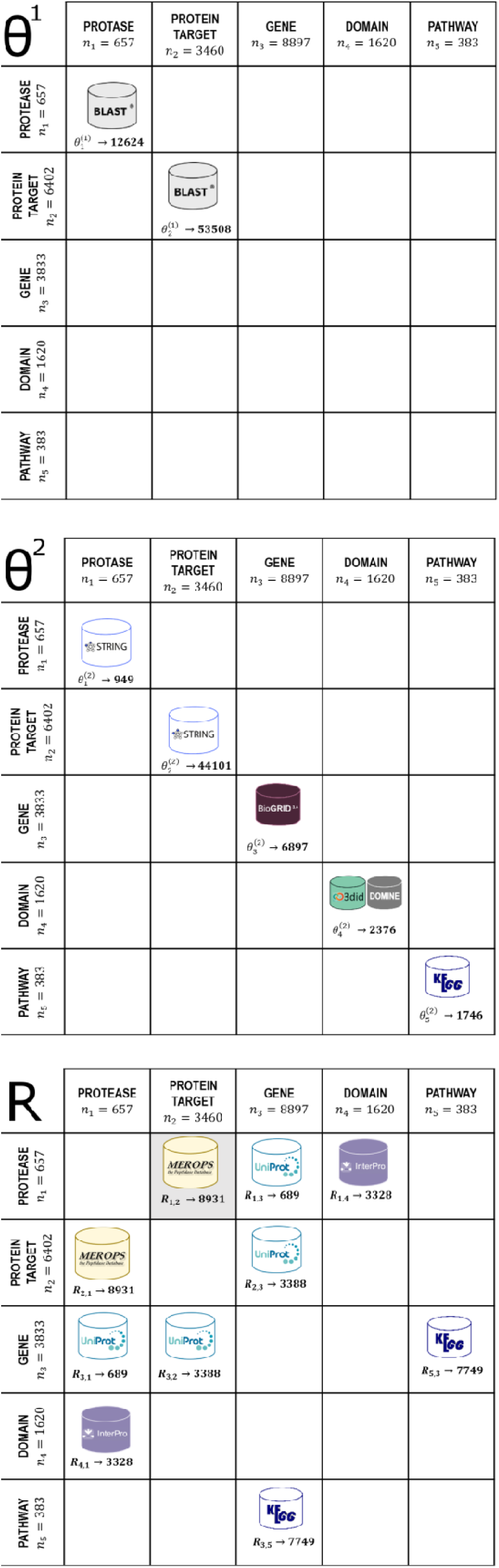
Data representation. Relational data are structured into sparse, block matrices and (same- and different-type data, respectively). We report the data types and sources.

### 2.2 R matrices: different data-type relations

Input matrices *R*_*ij*_ are then combined into a block matrix *R* * *R* ^*N* × *N*^, with *N* = ∑_*i*_ *n*_*i*_, containing all the available associations of *r* data types:

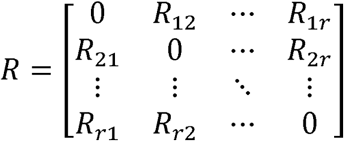

This matrix representation does not compress or alter the original structure of the data, which are simply juxtaposed in a block matrix, where a block is *R*_*ij*_. Note that *R* is a hollow matrix, being composed by input matrices only (i.e. same data types relations are not considered). Blocks in diagonal are empty. Also note that some blocks can be null due to missing (or meaningless) associations between the *i-th* and *j-th* data types. Furthermore, is typically very sparse, since the know relations are just a small fraction of all the possible combinations of data types. Values of input matrices must be bound to the [0, 1] interval, where 1 indicates the strongest association, while 0 represents the absence of relation or the lack of knowledge about it.

### 2.3 *θ* matrices: same data-type relations (constraints)

*θ*_*i*_ matrices are accounting for the data constraints in the tri-factorization algorithm, and record the associations between elements of the same kind (such as protease-protease or domain-domain relations). It can happen that multiple *θs* can describe constraint for the same data type, according to the following schema:

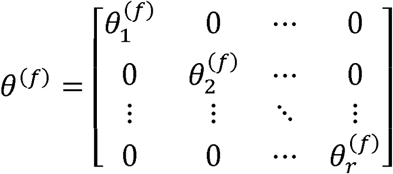

For example, in order to model the protein-protein relationships, we can use both a primary sequence similarity from BLAST and protein interactions described in STRING. In this way, we can derive two constraint (*θ*_S_) matrices to be used simultaneously. Being *f* the maximum cardinality of the,*θ*_*i*_matrices, we define *f* block diagonal, *θ(f)* matrices, with the same size of *R*.Each 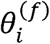 (if existing) is then the *i-th* block on a diagonal, and represents a data type or ontological entity, e.g., proteins, genes, pathways, etc. Constraint matrices are bound in the [-1,1] interval, where −1 indicates the strongest association, and 1 represents a negative association. We therefore can treat the negative values of *θ*s as must-link constraints; reversely, positive values of s indicate cannot-link constraints.

### 2.4 Data set

A first list of 657 human proteases cleaving 3460 human protein targets in 8931 interactions substantiated by experimental evidence has been obtained from MEROPS database. The protease-protein pair interactions represent the positive samples, while the non-interacting pairs represent the negative ones. STRING provided protein interactions data (threshold 0.7) for protease-protease and target-target associations, respectively 949 and 44410 pairs. Sequence similarity was measured with from BLAST, and filtered using a 10^−^10^^ e-value threshold. This procedure produced 12624 and 53508 elements for the protease-protease and protein target-protein target matrices, respectively. An InterPro analysis revealed 4723 domains expressed on 3328 protease-domain relations, and 13368 protein target-domain targets. 2376 domain-domain interactions were retrieved in Domine and 3did. Both proteases and protein targets have been mapped with UNIProt on their 3833 coding genes, as 689 protease-gene and 3388 protein target-gene relations emerged. Genes form 6897 interacting pairs on the BioGRID database. The genes expressing proteases and their targets map 7749 gene-pathway relations, as they are involved in 290 KEGG pathways. Pathways, in turn, form 1746 pathway-pathway relations. The assembled *R* and *θ* matrices are depicted in Figure 1.

### 2.5 Feature encoding by relational matrices

Both *R* are *θ* characterized by very high sparseness due to the fact that biological interactions are typically a tiny fraction of all potential interactions (Gilchrist *et al*., 2004; Orlowski *et al*., 2007). A target matrix *R*_*t*_ is selected among the block matrices. It will be reconstructed through tri-factorization into Ŕ_*t*_. The final goal is to learn the novel interactions, i.e. filling the gaps in t to unveil novel relations between the two specific data types of *R*_*t*_. The newly learned relations are to be found in the *dissimilarities* between *R*_*t*_ and Ŕ_*t*_ In our application, these data types are *proteases* and *protein targets*. This is obtained by factorizing the starting R matrix into the product of three terms. Each *R*_*i j*_block is tri-factorized 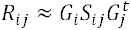. Here, *G* ∈ ℝ ^*Nxk*^ is a non-negative block diagonal matrix, and block *G*_*i*_ ∈ ℝ ^*n*_*i*_*xk*_*i*_^ represents the *i-th* data type, while *K* = ∑_*i*_ *k*_i_ *K*_i_ terms refer to the ranks, defining the dimension of the latent factors for the ***i-th*** data type. They are typically orders of magnitude smaller than the associated dimension (Žitnik and Zupan, 2015; Vitali *et al*., 2016). The objective function to be optimized is

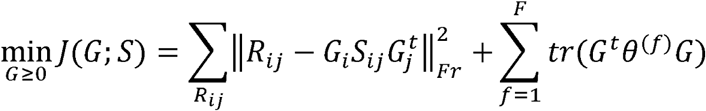

The first part of the objective function, 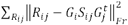 is the Frobenius norm of the difference between the original matrix and the tri-factorized one, and the second part of the objective function depends on the constraint matrices 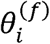 Note that without this second part, the objective function would simply lead to a tri-factorization without the contribution of the θs. The second term, 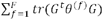, penalizes the cost function according to the must-link and cannot-link of the constraint matrices. Note that ***F*** represents the maximum multiplicity of *θ*, and, in our case ***F***=2 (Figure 1). The whole tri-factoriazion process is carried on through an iterative process detailed in the supplementary information. In summary, after initialization of the *G*_*i*_ factors, an alternate optimization *G* of *S* and is performed. So keeping *G fixed, S* is updated, and then keeping *S fixed, G* is updated. The update rules for the two types of matrices are obtained by computing the roots and the partial derivative of *J,* fixing the other matrix. Factorization ranks*.K*_i_ determine the number of columns in *G*. matri-ces, one per object type. Fixing the size of *G*, in other words, is a crucial factor for the data integration approach, as it determines the number of latent features condensing the information of each object type. (As reported in the Section *2.7*, we inferred the best factorization ranks according to a grid search cross-validation.) Stopping criteria is determined by either a maximum number of iterations, or two consecutive values *J*_*i*_, *J*_*i+1*_ such that *J*_*i*_, *J*_*i-1*_ <, where *T* is a fixed threshold (i.e., the variation between two iterations is small enough to consider the algorithm plateaued). Figure 2 depicts the core idea of feature representation and matrix reconstruction through tri-factorization.

**Fig. 2.**
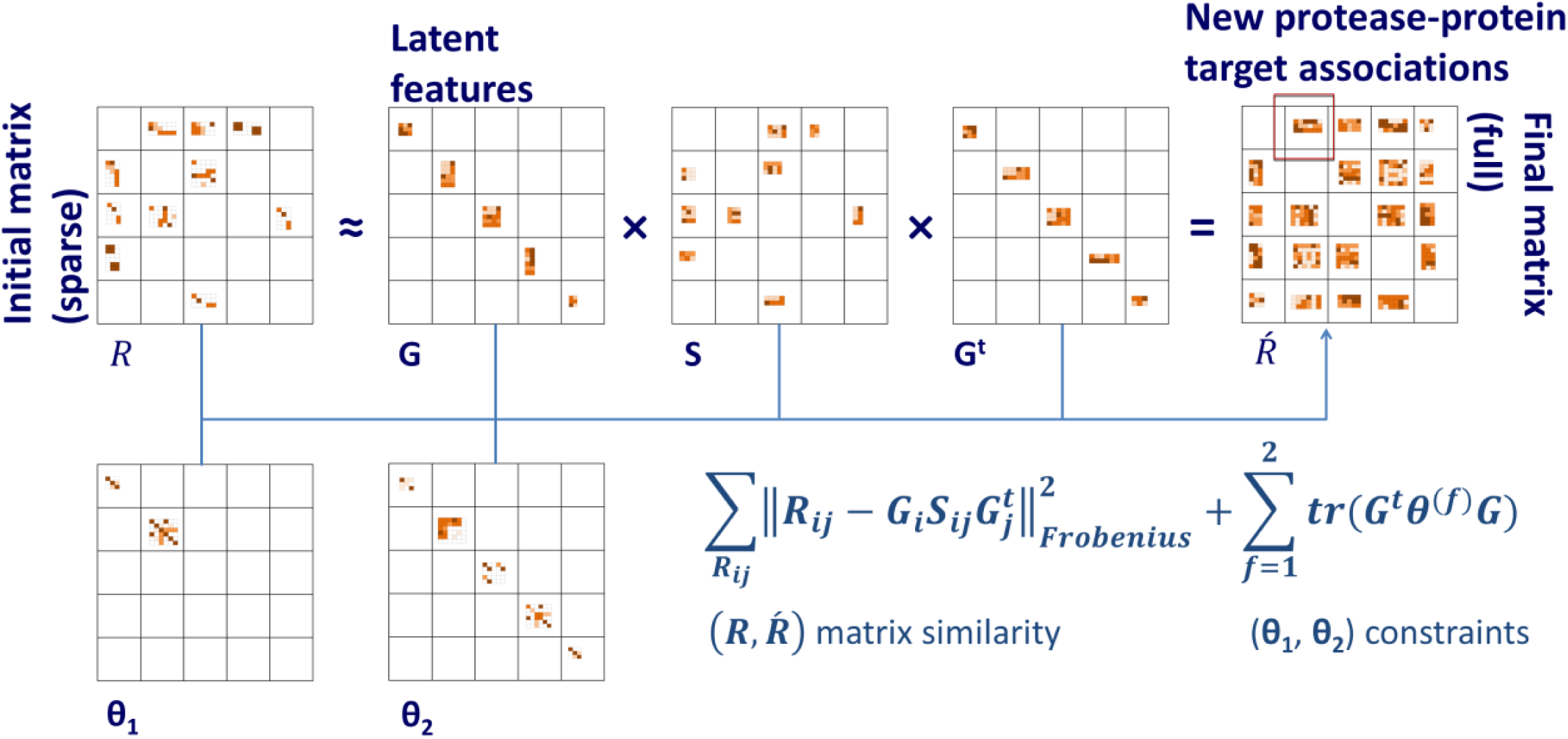
Data are embedded into three relational, sparse, same-size matrices:, and Each matrix is composed by concatenating smaller matrices describing the relation between objects (protease, protein targets, pathways, domains, and genes).is tri-factorized, and recomposed in by an iterative algorithm. This process both minimizes the dissimilarity between and (measured through the Frobenius norm) and forces to be compliant with the constraints imposed by and. Novel associations are found in the differences between and.

### 2.6 Inferring novel relations

Once Ŕ_*t*_ has been produced by the algorithm, a rule is needed to infer which of the newly Ŕ_*t*_ non-zero elements are to be considered newly predicted relations, i.e. predicted protease-protein target pairs. (Note that the matrix-assembling rule binding *R*_*t*_ to the [0, 1] interval does not apply to the tri-factorized Ŕ_*t*_.) Ŕ_*t*_ is used to compute a connectivity matrix *conn*_*m*_ a binary matrix derived from Ŕ_*t*_ where elements to be considered (*predicted* or *previously known*) relations are 1s, and unconnected elements are 0s. To set a row *r* and column *c* element *E*_*rc*_ of Ŕ_*t*_ as a newly predicted relation (a 1 in *Conn*_*m*_), its value should be higher than the average value of non-zero elements in a row *r* or column *c* of original target block matrix *R*_t_, as illustrated in Figure 3. Note that, since *R*_t_ here represents the proteaseprotein target relations, it is a binary matrix (cleavages are either present or absent/unknown, i.e., a protein is either a target or a not for each give protease). As consequence, the average value of *R*_t_ non-zero elements is always 1.

**Fig. 3.**
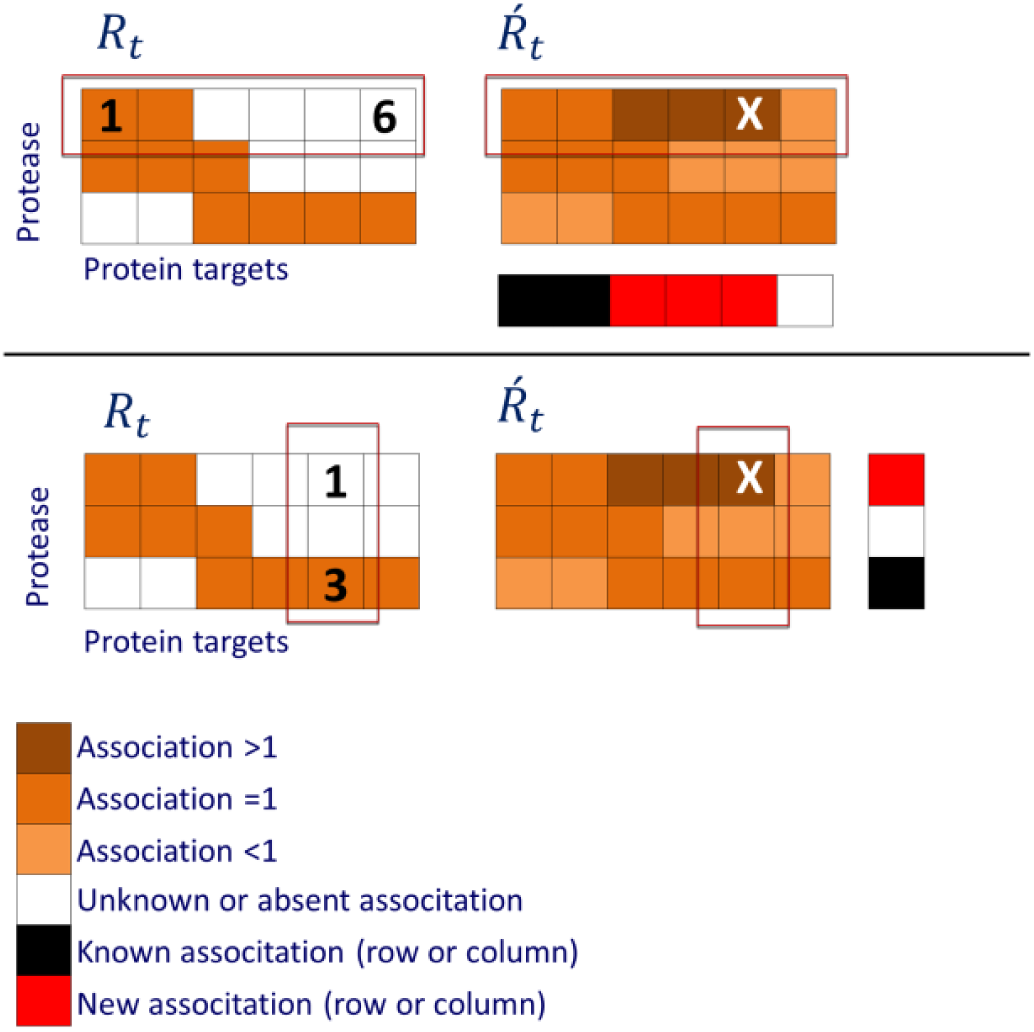
Discovering novel targets. This picture shows a toy example of the steps used to build a conne tivity matrix. Target reconstructed matrix is compared the original matrix by considering rows or columns. In, elements (targets) are either present (1, orange), or absent/unknown (0, white). In, elements can assume values higher (brown) or lower (light orange) than one. *Row criteria (top panel.)* If we consider the first row of, novel associations could appear in elements 3, 5 and 6, as elements 1 and 2 already indicate an existing association. Elements 3, 4 and 5 all show a value higher than 1, and can be considered as potential new target according to the row criteria. *Column criteria (bottom panel).* Let’s now show novel target from the column point of view. Considering the fifth column, we have one kno n interac-tion (element 3,). In the fifth column of, only element 1 shows a value higher than 1, and can be considered as a potential new target according to the column criteria. This element (marked by *x*) fulfills both its row and column rules, i.e. the putative new association is considered true for connectivity matrix if it satisfies both criteria.

Since the initialization of the algorithm is randomized, *nr* runs the algorithm will produce *nr* different *Conn*_*m*_ matrices. By summing all *Conn*_*m*_ s and divide it by the number of runs *nr* A final Consensus Matrix *C*_m_ is obtained, as the final output of the approach. Note that since the rule to populate, can be based on rows or columns, it is possible *Conn*_*m*_ to build both *C*_*m,row*_ ***ad*** *C*_*m,col*_

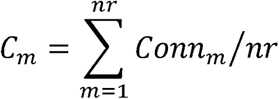

In other words, given a target *R*_t_ ∈ *R*^*n*_*i*_*×n*_*j*_^, *C*_*m*_ ∈ *Z*^*n*_*i*_*×n*_*j*_^ has elements bound to the [0, 1] interval. Each *C*_*m(i,j)*_ element reflects the number of times that relation has been predicted as positive in the *Conn*_*m*_s. For example, a relation, *C*_*m(i,j)*_ predicted as positive in half of the runs will have a value of 0.5; a relation never predicted as positive will have a value of 0;and a relation always predicted as positive will have a value of 1. Note that the output of our approach will therefore be a number in the interval [0, 1], solely denoting the likeliness or the absence/presence of a relation between the two elements of the target matrix. In our case, this relation represents the likeliness of a protein to be a target for a specific protease. As consequence, only the cleavage absence/presence is provided, and not the cleavage point. Not having an indication about the possible location of cleavage positions is the major limitation of our **spproach.**

### 2.7 Parameter tuning

We randomly split all the possible protease-protein target pairs into a training (85 %) and test (15 %) set, stratified on class. The considered parameters for this tri-factorization approach are (a) the stop criteria; (b) the number of runs *nr* to build *C*_*m*_; (c) the initialization; (d) the threshold(s) on *C*_*m*_ to call for a newly predicted relation; and (e) the factorization ranks *k*_*i*_ There is no clear consensus in literature about how to select these values (Žitnik and Zupan, 2015; Gligorijević *et al*., 2016). We therefore opted for an empirical grid search by cross-validation the training set, maximizing Matthews Correlation Coefficient (MCC), a suitable tool to measure performance of unbalanced class data sets such as ours. In order to ease the computational burden of the whole process, we fixed: the stopping criteria (a) by setting *T* = 10^−5^, according to literature (Žitnik and Zupan, 2015), and the maximum number of iterations to 10 thousand; the total number of runs for a single fold (b) was set to 5; initialization of G and S from random [0, 1] uniform distribution (c). Other parameters were optimized with a grid search. In particular, we set a double threshold to isolate newly pre-dicted) relations (d), searched among five thresholds [0.2, 0.4, 0.6, 0.8, 1] on both *c*_*m*_,_row_ ad a *C*_m_,_col_ factorization ranks were searched independently for each data type as 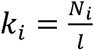, where *N*_*i*_ is the total number of non-zero elements of a data type *i* the overall associations modeled by all the related relation matrices *R*_*ij*_ and *R*_*ij*_ while. *l* is a scaling factor to be optimized in the interval [50, 100, 250, 500, 750, 1000]. Note that each data type *i* was independently considered for its scaling factor *l*. The grid search was performed with a 5-fold cross validation on the training set, measuring all the possible combination of parameters(d) and (e). Considered values for parameters are reported in the supplementary information.

## 3 Results

After the grid search optimization on the training set, the best parameter combination was *l*=500 for domains, genes, pathways, and proteases; and *l*=250 for protein targets. The thresholds for the Consensus Matrices were 0.2 for columns, and 0.2 for rows. We run our algorithm 10 times on the test set in order to compute the Consensus Matrices. On the test set, our approach with these specifics provided Specificity 0.999, Precision 0.88, ROC area 0.79, and MCC 0.446.

### 3.1 Comparison with other algorithms

We measured the performance of the proposed method against five well known other approaches: a BLAST search, Cascleave 2.0, PROSPER, PeptideCutter and PCSS. PoPS was considered as well, but we could not find a suitable implementation. Beside BLAST, all other algorithms have limited scope in terms of protease-target protein when compared to our more general approach. We therefore limited our comparison to the test set parts overlapping protease-protein target pairs predictable by the different algorithms. Table 1 reports details about these algorithm-specific test sets. Note that these algorithms provide multiple cleavage scores (i.e., the likely cleavage points along the primary structure of a target), while our approach outputs a dichotomy (i.e., the protein is or is not a target for a given protease), we could only compare ROC areas. For other algorithms, therefore, the highest cleavage score produced for a given target by a given protease was retained, rescaled in the [0, 1] interval, and used in the ROC. We run the models for BLAST, PROSPER and PeptideCutter; for PROSPER, Cascleave 2, and PCSS, we utilized the pre-computed scores provided by their respective websites. Some of the tested protease-protein target pairs, therefore, have very likely been included in the training sets of other algorithms. This condition is unfavorable for our approach, and implies results represent a best case scenario (i.e., an upper bound) for the predictions provided by other models. Our approach outperforms other algorithms, averaging a 17.8 higher ROC area. As consequence, our model seems to generalize better, to the point of providing better results even when compared on the Caspase-specific sets of Cascleave 2.0. Results are detailed in Table 1 and in Figure 4.

**Table 1.** Method comparison. Our approach outperforms other algorithms in the test subsets. Note that our algorithm is the only approach capable to provide predictions for all the available proteases. The subsets are obtained by considering the protease-protein target pairs of our vast test set that each algorithm is capable of classify.

| Subset | # of overlapping test-set pairs | ROC area |  |
| --- | --- | --- | --- |
|  |  | Other algorithm | Our method |
| BLAST | 340983 (all) | 0.65 | 0.79 |
| Cascleave | 2377 | 0.63 | 0.74 |
| PROSPER | 3574 | 0.51 | 0.77 |
| PeptideCutter | 1308 | 0.26 | 0.56 |
| PCSS | 2279 | 0.66 | 0.73 |

**Fig. 4.**
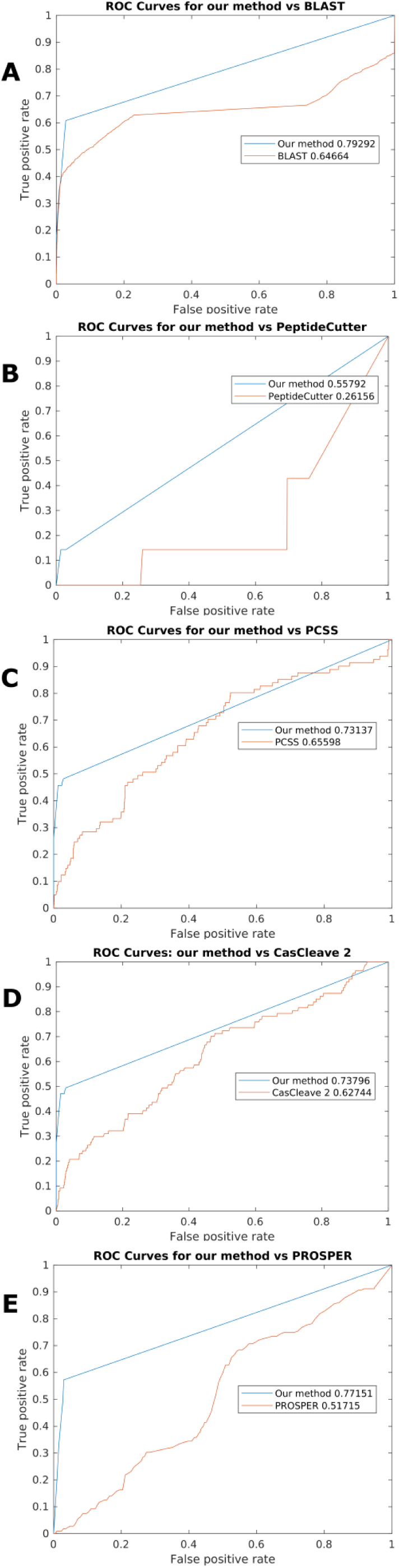
Comparison of our method vs other algorithms. Note that our approach does not provide a continuous score, but limited a set of 21 possible values indicating if a protein is (or is not) a target for a given protease. (This depend on the finite number of values an element can have in the ad Consensus Matrices, depending on the total number of runs. For this work, we opted for ten runs). Because of this, the great majority of our predictions correctly show a score of 0 (non-target), and the ROC c rve quickly becomes a straight line. Since other algorithms have a smaller prediction scope, in terms of protease families, we resolved to select all the predictions that overlapped with our test set pair, and calculate the area under to ROC curve. In order to compare the results, scores f om each algorithm were linearly rescaled between 0 and 1. (A) BLAST. The simple approach based on BLAST similarity is the only one able to generalize over *all* the ∼34100 combinations of proteases and candidate target proteins encompassed by our approach. (B) PeptideCutter. As reported in Table 2, there is only relatively small overlap (1308 pairs) between our test set and the capabilities of PeptideCu ter. Of these, only 7 are positive targets, which all return extremely low scores in PeptideCutter predictions. This, coupled with the tendency of PeptideCutter of providing high scores for false positive, justifies the particularly negative performance of the algorithm. (C) PCSS. (D) Cascleave 2.0 (E) PROSPER.

### 3.2 Inferring novel targets

After the comparison with other algorithms, we retained the best parameter set and applied our method not to the test set only, but to the whole protease-protein target matrix. In other words, we applied our method to all the collected data in order to infer novel proteaseprotein target relations, using 10 repetitions to build *C*_m_ (Section 2.6). We manually analyzed the best 54 scoring pairs, i.e., the ones fulfilling both row and column criteria 10 out 10 times. Twenty-five proteases are involved: one Cathepsin, two Calpains, two Caspases, thirteen Metalloendopeptidases, and seven Serine proteases. We found evidence of at least one pair confirmed as a real cleavage reported in literature, but not yet present in our MEROPS-derived data set, for all families. In particular, we found ten literature-confirmed cleavages: Cathepsin-D, Calpain-1, and Calpain-2 cleaving Angiotensiongen (Andrés, 2014; Jiang *et al*., 2008); Caspase-7 cleaving SREBF1 (Gibot *et al*., 2009); Matrix metallopeptidase-9 cleaving Decorin (Yang *et al*., 2014), CTGF (Hashimoto *et al*., 2002), and Prolargin (Zhen *et al*., 2008); t-plasminogen activator cleaving Complement C4-A (Barthel *et al*., 2012); ADAM28 cleaving FCER2 (Okada, 2017, 8); and Neprilisin-2 cleaving Neurotensin (Skidgel and Erdös, 2004). For all other pairs, we found (a) that the target is cleaved by another protease of the same family of the predicted one, and/or (b) that both the protease and the predicted target are involved in the same disease model, or the cleavage was inferred in a quantitative/qualitative mechanism, suggesting the plausibility of the cleavage event. These findings are summarized in Table 2, and provided in detail in the supplementary information.

**Table 2.** Candidate novel targets for proteases. Our approach retrieved 54 top scoring protease-protein target pairs, involving 25 proteases. Ten of the predicted targets are verified in literature. The rest are suggested as plausible by (a) having another protein target (or protease) of the same family being cleaved (or cleaving) the same pair element (Family); or (b) being involved in the same qualitative, quantitative or disease machinery (Mechanism).

| **Protease (Merops ID)** | **Verified** | **Family** | **Mechanism** |
| --- | --- | --- | --- |
| Cathepsin-D (A01.009) | 1 | - | 1 |
| Calpain-1 (C02.001) | 1 | - | 5 |
| Calpain-2 (C02.002) | 1 | - | 2 |
| Caspase-1 (C14.001) | - | 1 | - |
| Caspase-7 (C14.004) | 1 | 2 | - |
| MMP1 (M10.001) | - | 4 | - |
| MMP-8 (M10.002) | - | 1 | - |
| MMP-9 (M10.004) | 3 | 4 | - |
| MMP-3 (M10.005) | - | 2 | - |
| MMP-7 (M10.008) | - | 2 | - |
| MMP-13 (M10.013) | - | 1 | - |
| MMP-14 (M10.014) | - | 3 | - |
| ADAMTS4 (M12.221) | - | 1 | - |
| ADAMTS1 (M12.222) | - | 1 | - |
| ADAM28 (M12.224) | 1 | 2 | - |
| ADAMTS2 (M12.301) | - | 1 | - |
| ECE-1 (M13.002) | - | - | 1 |
| Neprilysin-2 (M13.008) | 1 | - | - |
| Chymase (S01.140) | - | - | 2 |
| Coag. factor Xa (S01.216) | - | - | 1 |
| PLAU (S01.231) | - | - | 1 |
| t-plasmin.activator (S01.232) | 1 | - | - |
| Plaminogen (S01.233) | - | - | 1 |
| Epitheliasin (S01.247) | - | - | 1 |
| Furin (S08.071)F | - | - | 2 |

## 4 Conclusions

In this paper we present a novel application, based on matrix tri-factorization of multiple, heterogeneous data sources, to infer novel protein targets for human proteases. The main limitation of the proposed algorithm is that the output consists on the presence/absence of the cleavage, without indication of the cleavage point. Another limitation is the lack of integration of third- and secondary structure data, as well as other physicochemical characteristics, such as the disordered regions, that are typically exploited by other algorithms. Both these limitations are implicit in the features encoding of our model, based on a matricial representation of ontological relations between several data types. We plan to integrate these features in a future work, which could exploit the presented method to infer the presence of the cleavage, and then apply another approach to estimate the cleavage site position along the primary structure. In contrast to previous works, focusing on single-protease models, our approach consists of a broad, general approach, for the first time encapsulating both protease-protein target knowledge and structured biological ontologies from biology in a single framework. Besides providing a larger scope, our model systematically outperforms state-of-the-art models.

## Acknowledgements

We are deeply grateful to (in random order) Sarah Boyd, Jiangning. Song, Francesco Pala, Neil Rawlings, and Jerico Revote for the invaluable help and support during the development of our work.

